# Loss of Vertically-Inherited Totiviruses and Toxin-Encoding Satellites in Killer Yeast Evidences Intracellular Conflict in Natural Populations

**DOI:** 10.1101/2025.05.03.652063

**Authors:** Thomas J. Travers-Cook, Sarah J. Knight, Soon Lee, Jana Jucker, Tamara Schlegel, Jukka Jokela, Claudia C. Buser

**Affiliations:** Institute of Integrative Biology, ETH Zürich, Zürich, Switzerland; Department of Aquatic Ecology, EAWAG, Dübendorf, Switzerland; Department of Zoology, University of British Columbia, Vancouver, Canada; School of Biological Sciences, University of Auckland, Auckland, New Zealand; Institute of STEM Education, St. Gallen University of Teacher Education, St. Gallen, Switzerland

**Keywords:** Mycoviruses, Infection, Inheritance, Interference competition

## Abstract

*Saccharomyces cerevisiae* is occasionally infected by totiviruses and their toxin-encoding satellites. Totiviruses and their satellites coexist but with an asymmetric dependence on the totivirus for maintenance inside the host cell. Satellites provide their yeast hosts with inhibitory toxins and the necessary self-immunity; loss of the satellite equates to loss of immunity. Because mycoviruses lack known extracellular stages, and sex is considered rare, mycoviruses are assumed to be transmitted vertically, implying infection states should correlate with host genotypes. However, totivirus–satellite coinfections are rarely examined in natural populations, leaving their associations with host genotypes poorly understood. We screened a multiyear population of *S. cerevisiae* isolates from New Zealand to examine the stability of host-virus associations over time, both within and across genotypes. While 55% of wild isolates harbored infections, only 37% of these included toxin-encoding satellites. Genotypes that persisted across years typically maintained consistent infection states. However, we observed stepwise transitions including acquisitions of totiviruses and satellites. Genotypes clustered strongly by infection state, supporting vertical transmission while suggesting that outcrossing is not responsible for the mycovirus acquisitions. Despite infection changes, genotype clustering by infection state remained intact, suggesting transitions are transient and that host genotypes may have optimal totivirus-satellite infection states.

## Introduction

Mycoviruses, those that infect fungi, tend not to cause overt harm or disease-like symptoms for their fungal hosts (Ghabrial et al., 2015). Mycoviral infections tend to be chronic and non-lytic, whereby transmission is generally restricted to vertical transmission from parent to progeny (Buivydaite et al., 2024; Ghabrial et al., 2015; Ghabrial & Suzuki, 2009). An intrinsic consequence of vertical transmission is selection for avirulence to prevent host mortality (Alizon et al., 2009; Anderson & May, 1982; Bull, 1994; Messenger et al., 1999). The fitness of both host and mycoviruses therefore tend to be coupled and enforced through partner fidelity feedbacks (Doebeli & Knowlton, 1998; Ewald, 1987; Herre et al., 1999; Sachs et al., 2011). Sexual reproduction can enable transmission of mycoviruses to new host lineages, although outcrossing is believed to be rare relative to the norm of clonal reproduction and intra-tetrad mating events (Bennett & Turgeon, 2016; Nieuwenhuis & James, 2016). Mycoviruses are therefore believed to diverge with host genotypes, which is supported by observations over macroevolutionary time (Goker et al., 2011).

*Saccharomyces cerevisiae* often harbours mycoviruses. These extrachromosomal genetic elements include both viral dsRNAs and plasmid DNAs (Wickner, 1996a; Wickner et al., 2013). The most well-studied of these is the monopartite dsRNA totivirus (family Totiviridae), which on occasion is found to coinfect *S. cerevisiae* with a separately encapsidated monopartite dsRNA satellite nucleic acid (Magliani et al., 1997; Schmitt & Breinig, 2002, 2006). Totiviruses are believed to reside permanently in the host cell, and replicate autonomously, due to their possession of two overlapping open reading frames (ORFs) enabling encapsidation and replication (Dinman et al., 1991; Dinman & Wickner, 1994; Fujimura et al., 1992; Icho & Wickner, 1989). Satellite nucleic acids of totiviruses are obligatorily dependent on the machinery and resources of totiviruses (Bostian et al., 1980b). However, the satellite’s single ORF encodes a preprotoxin – an unprocessed precursor of a mature, secretable toxin – which also encodes functional immunity for the host (Dignard et al., 1991; Hanes et al., 1986; Schmitt & Tipper, 1995).

The secreted killer toxin can be lethal to competitors of the host strain that do not possess the same totivirus-satellite coinfection type or have evolved resistance (Hanes et al., 1986). Satellite-encoded toxins are generally believed to be used by *S. cerevisiae* as an interference competition strategy for invading novel environments or for preventing invasions by distantly related genotypes lacking the coinfection (Boynton, 2019; Travers-Cook et al., 2023). It has also been proposed that satellite-encoded toxin production acts analogously to toxin-antitoxin systems in bacteria (Jurenas et al., 2022; Kast et al., 2015; Wickner & Edskes, 2015), whereby loss of the satellite nucleic acid leads to immediate competitive exclusion by clonal neighbours that maintain their toxin-encoding coinfection.

The totivirus-satellite association may be best considered under the context-dependent mutualism framework (Bronstein, 1994). There is an asymmetric dependence of the satellite on the totivirus. Unlike many mycoviruses, the totivirus detriments host fitness by inducing proteostatic stress at high temperature under laboratory conditions (Chau et al., 2023a). The satellite hijacks capsid proteins and polymerases for its own replication, which may slow the totivirus replication rate or encourage selection for virulent mutants of the totivirus (Bostian et al., 1980b; Taylor et al., 2014). While there may be selection on totiviruses to suppress resource hijacking by the satellite, removal of the satellite can result in host toxin sensitivity and mortality by the actions of clonal neighbours (Marquina et al., 2002; Woods & Bevan, 1968). Depending on density-dependence and the frequency of toxin-based interference competition, this may prevent yeast without satellite infections from fixating (Wickner & Edskes, 2015). *S. cerevisiae* has mechanisms in place to suppress totivirus proliferation, which are most effective against coevolved totiviruses whom they have adaptations in place for (Gao et al., 2019b; Rowley et al., 2016). *S. cerevisiae* can be cured of the satellite with relative ease under high temperature treatments (Wickner, 1974). The bias of vertical transmission of totivirus-satellite coinfections has selected for them to have minimal impact on host gene expression, demonstrated during artificial infection-curing experiments (Luksa et al., 2017; McBride et al., 2013).

It is not known whether infections are stable and consistent within host genotypes in natural populations of *S. cerevisiae*. Beyond cytoplasmic conflict between totiviruses and their satellites, rare sexual events may mix host-virus-satellite lineages and perhaps result in progeny having different mycoviral infections than those of their parents. Though sexual reproduction is considered to infrequently occur (Bennett & Turgeon, 2016; Nieuwenhuis & James, 2016), it is the suspected explanation for the occasional mismatching of totivirus-satellite combinations from their native pairings (Quintero-Blanco et al., 2022). Killer phenotype bearing strains from broad geographic origins have been found to be somewhat phylogenetically clustered within strains without this phenotype, yet killer phenotypes are evidently gained and lost (Pieczynska et al., 2013). Yeast isolates from the same sampling sites over time have rarely been screened for totivirus-satellite coinfections (Starmer et al., 1987), such that the frequency, or infrequency, of gains and losses of *S. cerevisiae*’s mycoviruses in wild populations is not well studied. It is undetermined whether infection states diverge with their host genotypes, or whether host-virus-satellite conflict and sexual reproduction are processes capable of distorting associations between host genotype and infection status.

In this paper, we utilise a multi-year collection of co-occurring vineyard-associated *S. cerevisiae* isolates from across New Zealand covering years 2018, 2019 and 2021, to explore the prevalence and persistence of totivirus-satellite (co)infections in natural populations of *S. cerevisiae*. To do so, we quantify the prevalence of the different infection states in this vineyard-associated population and co-divergence of infection state with the host genotype. We also examine the frequency of genotype-level transitions in infection state and whether these transitions disrupt associations between host genotypes and infection status.

## Materials and Methods

### Yeast Sample Isolation

We studied a collection of *S. cerevisiae* isolates from vineyards across the Hawke’s Bay and Marlborough regions of New Zealand. Yeast isolates were sampled in 2018, 2019 and 2021 from a set of 25 vineyards (Table S1). Grape varieties at each vineyard belonged to either Merlot, Pinot Noir or Sauvignon Blanc (Table S1).

Fruit was collected one to three days prior to the commercial harvest. For Sauvignon blanc, whole bunches were hand harvested from vines using sterile snips from nine random locations within each vineyard block, totaling 40 kg of fruit from each vineyard site. For each vineyard site, the fruit was crushed and destemmed mechanically under carbon dioxide (CO2) cover and 30 ppm of SO2 (Potassium Metabisulfite) was added to the must. The must was chilled at 6 °C for 1 hour on skins before being pressed in a balloon press at 2 bar for 3 minutes and 4 bar for 14 minutes. The pressed juice was left to cold settle overnight at 6 °C before being racked off lees under CO_2_ cover into 18 L fermentation vessels. Lees were added back to adjust turbidity to between 150-200 nephelometric turbidity units (NTU). The juice was warmed to 16 °C for uninoculated fermentation. Fermentation was performed at and monitored by weight loss (El Haloui et al., 1988). Once fermentation began (weight loss of >100 g in 24 h) a standard complex nutrient addition of 600 ppm (0.6 g/L) Nutristart® (Laffort®, Bordeaux, France) was added. Fermentation was considered complete when no weight was lost for two consecutive days and the residual sugar concentration was < 2 g/L as determined by AimTab™ reducing substances tablets (Germaine Laboratories Inc., San Antonio, TX, USA). Post-fermentation, a 1 mL sample was taken and stored in 15 % glycerol at -80 °C whilst awaiting processing. All equipment was washed with water and sanitized with 70% ethanol between samples.

For red wines (Merlot and Pinot Noir), whole bunches were hand harvested in the same manner to a total of 17 kg per site. The fruit was destemmed by hand under CO_2_ cover and 15 ppm of SO_2_ (PMS) was added to the must. The must was cold soaked at 6 °C for three days and the cap (grape skins, pulp and stems) was submerged twice during this time. If the pH was >3.6, tartaric acid was added to drop it below this level. It was then warmed to 25 °C for uninoculated fermentation and monitored during fermentation to ensure the temperature peaked between 28-32 °C. As with the Sauvignon Blanc fermentation, 600 ppm (0.6 g / L) Nutristart® (Laffort®, Bordeaux, France) was added as fermentation began. During the course of fermentation, the cap was manually plunged once daily and it was deemed complete once the sugar level dropped below 2 g / L as determined by AimTab™ reducing substances tablet (Germaine Laboratories Inc., San Antonio, TX, USA). A 1 mL sample for microbial analysis was taken post-fermentation and before the wines were pressed from each wine and stored in 15% glycerol for later analysis.

### Identification of Saccharomyces cerevisiae

Ferment samples were serially diluted onto standard YPD agar plates (1 % yeast extract, 2 % peptone, 2 % dextrose, 2 % agar powder) and incubated for 2 days at 20°C. Afterwards, 96 colonies were picked from each sample for further analysis. Genomic DNA was extracted by incubating colonies at 37 °C for 30 minutes in a pH 7.5 20 µL solution of 1.25 mg/mL^-1^ zymolyase dissolved in 1.2 M sorbitol and 0.1 M KH_2_PO_4_ followed by 10 minutes of incubation at 95 °C. A multiplex PCR was used to confirm that the yeast isolates belonged to *S. cerevisiae* or *S. uvarum* (de Melo Pereira et al., 2010). The SacScreen PCR reaction contained 1x KAPA 2G Readymix (Kapa Biosystems, Wilmington, MA, USA), 0.4 µmol/L^-1^ of each SacScreen primer (Pereira et al., 2010), ∼10 ng DNA template and H_2_O to a total volume of 10 µL per reaction. The SacScreen PCR settings were as follows: an initial denaturing step of 95 °C for 3 minutes, followed by 35 cycles of denaturation (94 °C for 30 seconds), annealing (55 °C for 30 seconds) and extension (72 °C for 2 minutes), all of which was concluded with a final extension step at 72 °C for 5 minutes (Nadai et al., 2018). Products of the SacScreen PCR were visualized with gel electrophoresis to confirm membership as either *S. cerevisiae* or *S. uvarum*, all of which were *S. cerevisiae*.

### Microsatellite Genotyping and Genotype Assignment

Microsatellite markers were used to genotype eight colonies identified as *S. cerevisiae* per vineyard-year combination. A multiplex PCR was conducted for each *S. cerevisiae* colony, of which 10 µL reactions were made up of 5 µL QIAGEN Multiplex mastermix (Venlo, Netherlands), 1 µL of a 20-primer mixture (forward and reverse primers for 9 informative microsatellite loci as well as primers for mating types), 2 µL genomic DNA and 2 µL H_2_O (Richards et al., 2009b). The PCR settings involved an initial denaturation step at 95 °C for 15 minutes, followed by 35 cycles of denaturation (94 °C for 30 seconds), annealing (55 °C for 30 seconds) and extension (72 °C for 2 minutes), concluded by a lengthened extension period (60 °C for 6 minutes). PCR products were genotyped at Auckland Genomics on an Applied Biosystems (Waltham, Massachusetts, USA) 3130XL Genetic Analyzer. The ensuing trace files were analyzed using the Geneious bioinformatics platform (Biomatters, Auckland, New Zealand). Of the 9 microsatellite loci that isolates were initially scored for, 8 are used in the analyses hereafter (Table S2). The YBR240C marker was excluded because it often had too many peaks on the trace files.

Much of our analyses described hereafter were conducted with a refined dataset with complete microsatellite data because some of the methods used here do not handle missing data well. We designed a conservative pipeline to assign genotypes with missing data at microsatellite loci to genotypes with more complete microsatellite profiles. We first assigned genotypes for those with complete microsatellite profiles over the eight microsatellite loci, then we iteratively assigned the other isolates to these genotypes based on whether their available data could be nested into the microsatellite profile of only one other genotype. If an isolate with missing microsatellite data could be assigned to multiple other more complete genotypes based on its available microsatellite data, it was considered to have an ambiguous genotype assignment and discarded from analyses that utilised genotype assignment or microsatellite information (Discriminant Analysis of Principal Components and Markov Chain Transition Probabilities). If the isolate’s available microsatellite profile was not a complete match with any of the previously defined genotypes, it was treated as a unique genotype, but again removed from analyses (Discriminant Analysis of Principal Components and Markov Chain Transition Probabilities). Whilst this approach resulted in a number of isolates with ambiguous genotype assignments being excluded from analysis (14.9%), we could avoid inflating the number of multilocus genotypes that were present due to missing data, whilst maintaining the genotype distribution. We established a genotype accumulation curve for the refined population without missing data and visually compared this to genotype accumulation curve achieved when including isolates with missing data. Each locus was randomly sampled 10000 times to create the distribution.

### dsRNA Extraction and Virus Typing

To establish the viral infection state of each yeast isolate, dsRNA was extracted from each yeast isolate using a modified version of Crabtree et al. (Crabtree et al., 2019), in which only a single wash buffer (1x STE; 16% EtOH) step was applied, as this could prevent the unnecessary loss of dsRNA. The dsRNA precipitate was dissolved in molecular grade water before visualisation using agarose gel electrophoresis. The presence of the totivirus in the yeast isolates was confirmed by a band at ∼4.6-4.8 kb, whilst the presence of the toxin-encoding satellite was confirmed by a band at ∼1.6 kb. Where an absence of dsRNA bands was found across a whole set of extractions, extraction sets were repeated to eliminate the possibility of erroneous extractions being responsible for the observation of virus-free isolates. From this, each yeast isolate could be allocated to one of three infection state groups: “Totivirus-satellite coinfections”, “Totivirus infections” and “Infection-free”. These groups, defined by the numbers of bands observed from the gel electrophoresis, are unambiguous and discrete (Image S1). These three groups capture all possible combination as satellites can only inhabit yeast when the totivirus accompanies it, and as expected, it was never observed to infect alone. We used a chi-squared test to determine whether the observed counts of each infection type significantly differed from the expected equal distribution.

### Temporal Changes in the Proportions of Different Infection States

To assess the effect of the *year* on the relative proportions of each infection state found, compositional analysis was conducted using a multinomial Dirichlet regression model (Douma & Weedon, 2019). Within vineyards there were instances of infection types being absent. To overcome this technical problem for calculating test statistics we added a small value of 1 x 10 ^-5^ to each infection count before calculating the proportions. This transformation ensures that the proportions derived from the count data fit within the [0,1] limits of the Dirichlet regression model. The same value was added to the denominator as a normalisation factor to ensure all proportions were within the [0,1] limits. In the model, *year* was treated as a fixed effect and *vineyard* as a random effect, utilising the Dirichlet family to account for the compositional nature of the data. The model was implemented using the brms package in R (Bürkner, 2017), employing 10 Monte Carlo Markov chains (MCMCs) of 5000 iterations for parameter estimation, with the first 2500 iterations serving as a burn-in period. A total of 67 vineyard x year combinations were analysed, accounting for variability amongst 25 vineyards.

### Intra-Genotypic Infection State Transition Probabilities

To explore the consistency of coinfections within individual genotypes over time, a discrete-time Markov chain (DTMC) was constructed. This excluded individuals with ambiguous genotype assignments (based on missing microsatellite data), and genotypes that were only observed in one year, resulting in a total of 28 genotypes with multi-year presences (24.4 %) being used for the calculations of transition probabilities. Transition matrices were created for each genotype, capturing the changes in infection state from one year to the next. Transition probabilities were computed based on the prevalence of each genotype in the current and subsequent years. Transition matrices were then aggregated across genotypes and years and normalized to ensure that each row sums to one, making the probabilities comparable across infection states. Confidence intervals for the transition probabilities were calculated across data for individual genotype transition probabilities across pairs of states. All analyses were conducted using the R programming language (R Core Team. 2023), and plotted with ggplot2 (Wickham, 2011).

### Discriminant Analysis of Principal Components (DAPC)

Discriminant analysis of principal components (DAPC) was used to cluster genetically similar individuals according to their multilocus microsatellite profiles (Jombart et al., 2010). DAPC reduces the dimensionality of the microsatellite dataset with principal component analysis, to which discriminant analysis partitions the between- and within-group variation component to maximise the between-group differences. Discriminant analysis therefore summarises the between group genetic differentiation. We reduced the dataset to isolates that have complete microsatellite genotype at the eight microsatellite loci used, after our genotype assignment method described previously.

Three DAPC models were calculated to untangle the contributions of genotype frequency and variable infections within genotypes on how genotypes cluster by infection state, all with cross-validation to determine the number of principle components (PCs) that achieve the highest mean successful assignment (MSA) and lowest mean squared error (MSE). We first conducted the DAPC with all individuals with complete microsatellite profiles (n = 414), with infection state set as the prior; this approach makes genotype weighting proportional to their frequencies. For this model, retaining 26 PCs achieved the highest MSA (0.861) and lowest MSE (0.153). Next, we applied a clone correction to remove replicate isolates of the same genotype. The first clone correction of the population included all genotypes and their replicates with different infection states (n = 98); this method gives additional weight to genotypes that have variable infection states and was used to see how infection state transitions disrupt the MSA. The highest MSA (0.728) and lowest MSE (0.309) was found when 10 PCs were retained. Finally, we further refined the collection to include only genotypes that were not observed to have variable infection states across the collection (n = 67). The final approach removes the weighting bias introduced by genotypes that have multiple infection states across the collection and explores only genotypes with consistent infection states. This model had the highest MSA (0.902) and lowest MSE (0.162) when 12 PCs were retained.

## Results

In total, 536 diploid yeast colony isolates were microsatellite genotyped across 25 vineyards over one to three years. Prior to any form of genotype correction, 188 genotypes were identified. Correcting for ambiguous assignment by nesting isolates with missing data into more complete genotypes as described in the methods section, resulted in 119 genotypes and 80 individual isolates with ambiguous microsatellite genotype assignments (14.9%) – these isolates were excluded from analyses that required genotype information. Isolates with complete microsatellite profiles (eight loci) after our assignment method totaled to 414 individuals of 80 genotypes (77.2% of individual isolates). The decline from 119 to 80 genotypes is due to the presence of individuals with unique microsatellite profiles yet with missing data. The genotype accumulation curve demonstrates that the asymptote has been mostly reached in the number of loci needed to discriminate the individuals into genotypes (Figure S1).

Of the 536 isolates in the collection, we successfully established virus profiles for 534 individuals (99.6%). In summary, 240 were free of any form of cytosolic dsRNA (infection-free), 185 isolates possessed a gel band corresponding to the sole presence of totiviruses (totivirus infection) and 109 isolates possessed gel bands corresponding to the presence of the totivirus and its coinfecting satellite (totivirus-satellite coinfections; Figure 1A). Less than half of isolates with the totivirus infection were observed to also have a toxin-encoding satellite (37.1%). Using a chi-square test, we determined that the observed prevalence of each infection type were significantly different from the expected equal distribution (χ^2^ = 48.62, df = 2, p < 0.001). Of the 119 MLGs identified, 28 MLGs appear in multiple years and 15 were found to have multiple viral profiles (Figure S2).

**Figure 1.**
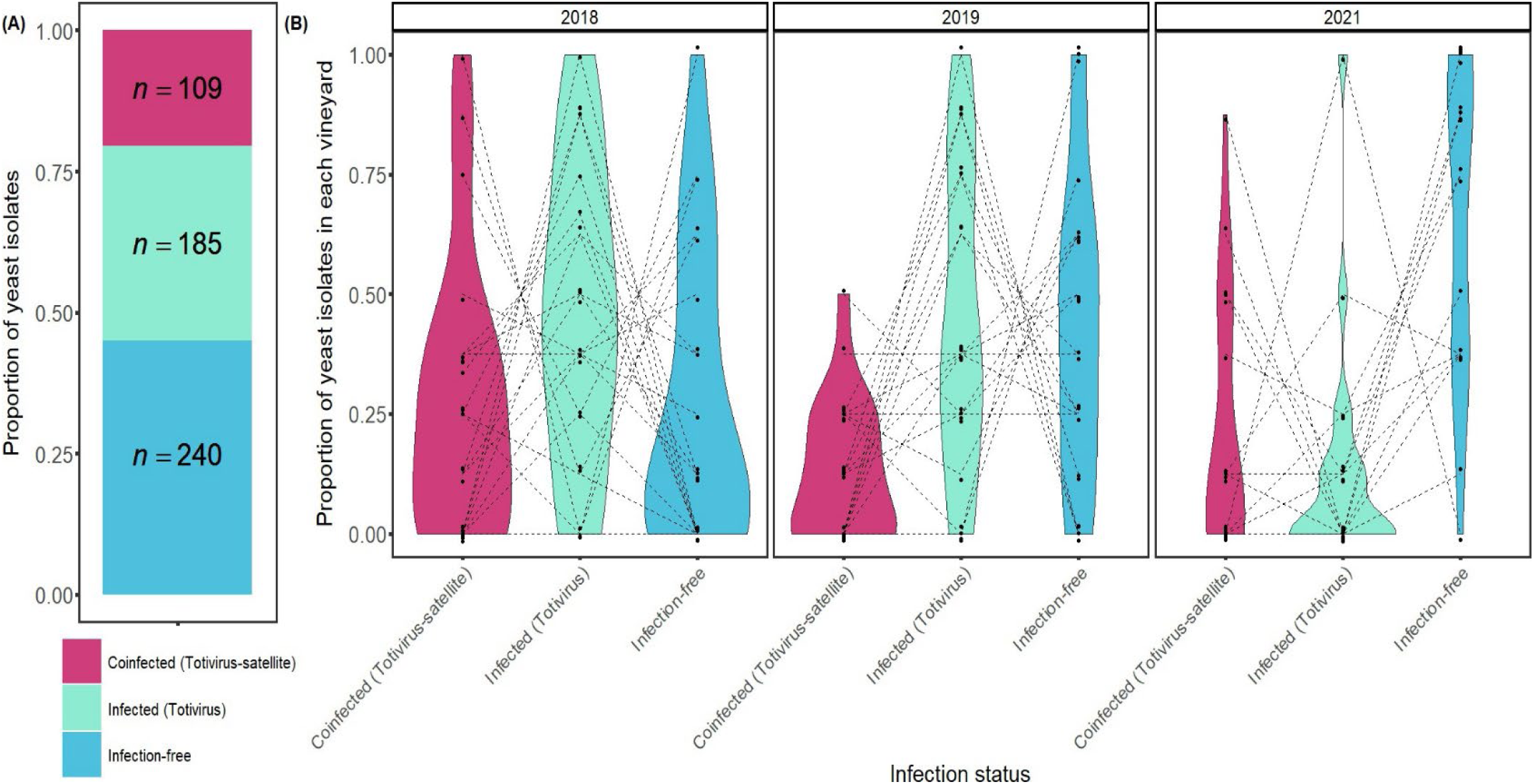
The proportions of each infection status in the population. (A) The overall proportions of each infection status in the population with the frequency (*n*) of each seen in each stack of the bar plot. (B) The proportions of each infection status across year in individual vineyards. Dotted lines connect proportions from the same vineyards and demonstrate that the proportions equal to 1.

### Temporal dynamics of yeast with different infection states

A multinomial Dirichlet regression model was employed to assess the effect of year on the relative proportions of different infection statuses across vineyards, with the infection-free group serving as the reference. The random effect of vineyard revealed moderate variability in infection type proportions across sites, with the standard deviation of the intercept for totivirus infections estimated at 0.33 (95% CI: [0.01, 0.81]) and for totivirus-satellite coinfections at 0.21 (95% CI: [0.01, 0.58]). This suggests greater vineyard-level variability in the prevalence of totivirus infections compared to totivirus-satellite coinfections. Relative to the reference year of 2018, totivirus infections had a moderately lower prevalence in 2019 (estimate: -0.60; 95% CI: [-1.39, 0.19]; Figure 1b) and a substantially lower prevalence in 2021 (estimate: -1.88; 95% CI: [-2.72, -1.05]; Figure 1b). Totivirus-satellite coinfections were also less prevalent in 2019 (estimate: -0.74; 95% CI: [-1.55, 0.09]; Figure 1b) and in 2021 (estimate: -1.19; 95% CI: [-2.02, -0.36]; Figure 1b).

### Intra-genotypic transition probabilities

The genotype infection state transition probability matrix indicates that if the same genotype was found in multiple years, it most likely maintained its infection status (diagonal, Figure 2). There were, however, rare transitions to all of the other infection states (off-diagonal; Figure 2). The transition matrices suggest that state transitions occur in a stepwise manner. Transition probabilities from coinfection to infection-free, and vice versa, were rarely observed, if at all (0-0.01; Figure 2). Maintenance of a totivirus infection was most probable of all (0.822), and remaining infection-free was the most improbable (0.717) among the non-transitions. Nonetheless, transitions between all states were observed at least once, including gains of totiviruses and/or satellites (Figure 2).

**Figure 2.**
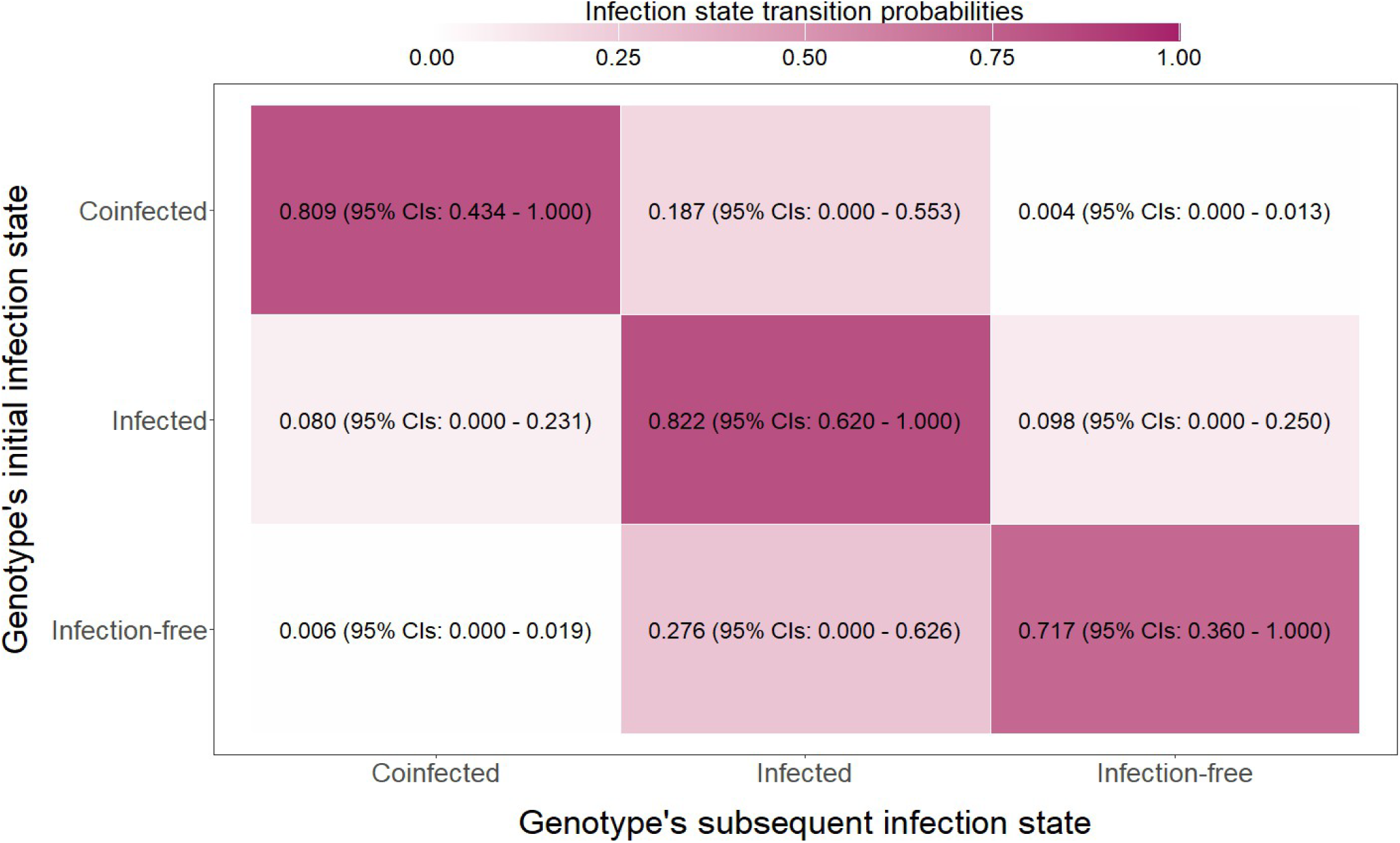
Transition probabilities across the infection states. Values in the center of each cell constitute the average transition probability across all genotypes, as well as the 95% confidence intervals. Each row sums to 1; however rounding to three decimal places introduces slight discrepancies.

### Discriminant analysis of principal components (DAPC)

To visualise how infection state interrelates with *S. cerevisiae* genotype, the results of the two linear discriminants of the DAPC were plotted with infection state as a prior, with the three previously defined approaches (Figure 3). Using all individuals with full microsatellite profiles after our customized genotype assignment (Figure 3a), we see strict clustering of individuals by their infection states. The first linear discriminant alone was capable of near-perfect discrimination of uninfected individuals from those with totivirus-satellite coinfections. The second discriminant was equally capable of discriminating totivirus-infected individuals from the other two groups. Infected isolates are at an intermediate level between coinfected isolates and uninfected isolates on the first linear discriminant. The coinfection virus profile is the most distinct of the three infection states, whilst infected and uninfected isolates cluster more frequently with each other.

**Figure 3.**
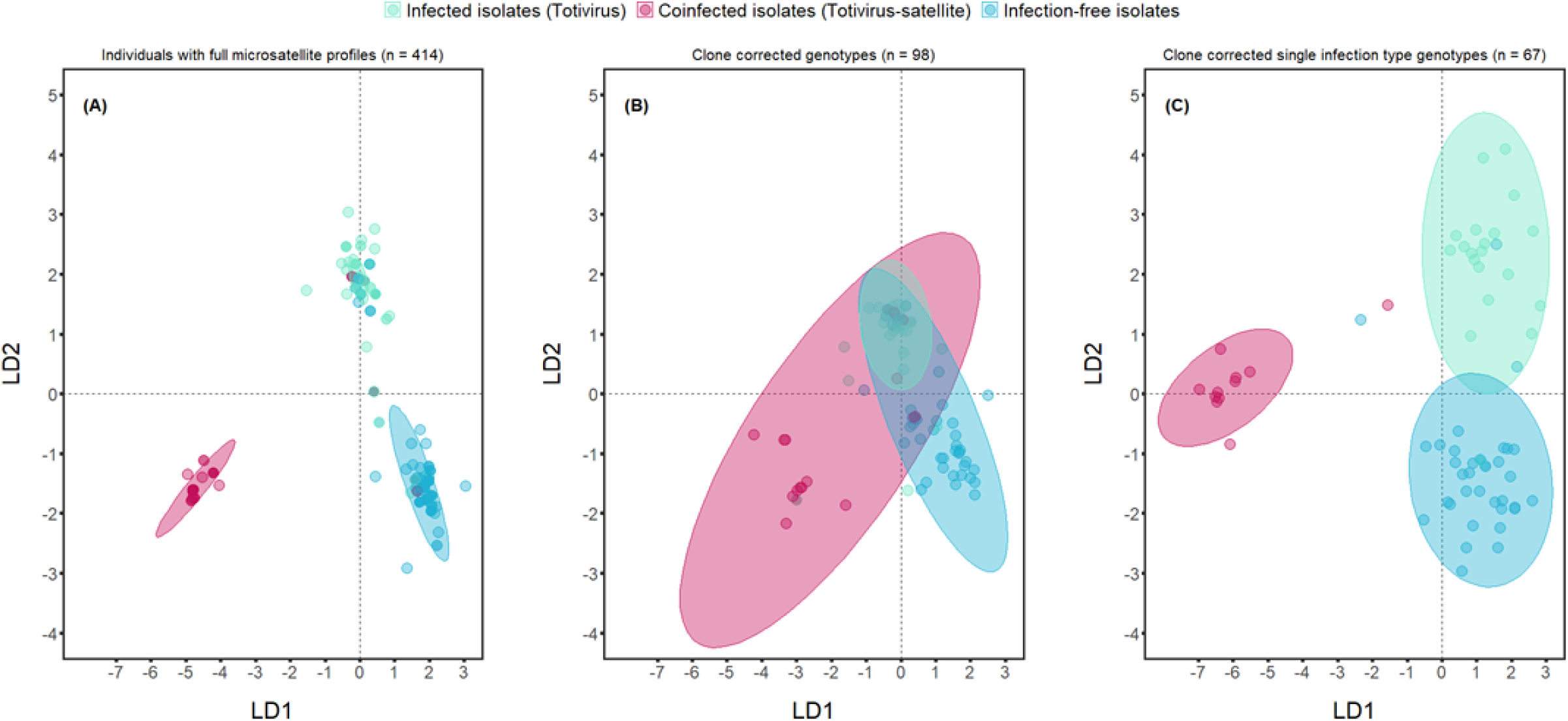
The two linear discriminants of the discriminant analyses of principal components (DAPC) analysis using microsatellite data with viral infection status as a prior. Circles represent the individual isolates and their colour indicates the individual’s infection status, whilst the infection status prior genotype space with 95% confidence intervals are represented by inertia ellipses. (**A**) All individual isolates that have complete microsatellite profiles after application of our conservative genotype assignment method (n = 414); (**B**) Clone corrected genotypes based on all individuals isolate that have complete microsatellite profiles, with replicates of genotypes with multiple infection states (n = 98); (**C**) Clone-corrected genotypes based on all individual’s isolate that have complete microsatellite profiles, but excluding genotypes that are observed to have isolates with different infection statuses (n = 67).

This distribution of the various infection states over the two linear discriminants is also observed across the other two models, but with noticeable differences (Figures 3b and 3c). For the clonally corrected genotypes including those with multiple infection states (Figure 3b), all three infection state priors coalesce and overlap in the DAPC coordinate space of the infected isolates, such that the inertia ellipses of the coinfected and uninfected prior groups extend towards 0 and past it (towards negative values for uninfected group and towards positive values for the coinfected group) on the first linear discriminant and toward positive values on linear discriminant 2. Once the population is further refined to remove genotypes that were observed to have multiple infection states (Figure 3c), we see the reemergence of distinct clusters, with some overlap between the totivirus-infected and uninfected isolate clusters. The coinfected prior grouping is again isolated, though one genotype evidently clusters better with the infected isolates.

## Discussion

*S. cerevisiae* is often hosting the autonomous dsRNA totivirus and its non-autonomous dsRNA satellite that when together encode the toxin-producing killer phenotype of *S. cerevisiae*, as well as the associated immunity (Schmitt & Breinig, 2002, 2006). This phenomenon is well-studied under laboratory conditions (Dignard et al., 1991; Hanes et al., 1986; Marquina et al., 2002; Wickner, 1996a), yet the prevalence of totivirus-satellite coinfections in natural populations is less well studied (Boynton, 2019; Chang et al., 2015; Starmer et al., 1987; Travers-Cook et al., 2023). It remains unclear whether totivirus-induced proteostatic stress in the host (Chau et al., 2023), the intrinsic conflict of asymmetric dependence of satellites on totiviruses, and host sexual reproduction, disrupt what should be, based on present knowledge, a strictly vertically transmitted mycoviral infection (Quintero-Blanco et al., 2022; Taylor et al., 2014; Travers-Cook et al., 2023).

We found that approximately half of the collection was infected by a dsRNA totivirus (55.1 %), whilst toxin-encoding satellites were only found to coinfect with totiviruses in a third of these isolates (37.1 %). The proportion of isolates infected by totiviruses in this collection was similar to other screening studies, however coinfections were found to be considerably lower in frequency when compared to the earlier studies (Crabtree et al., 2023). The proportion of yeast isolates with coinfections was found to be highly variable both across vineyards within years and between years (Figure 1b). Such variability across vineyards in representation of each infection state demonstrates a discordance between local and regional processes governing the presence of each infection type. Yeast isolates with each of the infection states were found to represent whole vineyard-isolate sets (Figure 1b).

Yeast isolates with totivirus-satellite coinfections experienced a major decline from 2018 into 2019, which was largely due to a single genotype (Figure S2) experiencing local extinctions across many of the vineyards it was previously present in, as well as a near-complete extinction across all vineyards. The drivers of this genotype’s disappearance are unknown, though we hypothesize that the redundancy of toxin production, when resistance emerges, may be involved (Boynton, 2019; Travers-Cook et al., 2023). The overall relative rarity of yeast isolates harboring the competitively advantageous toxin-encoding satellites suggest that negative frequency-dependent selection regulates their prevalence, thereby maintaining the infection-state polymorphism in the population (Christie & McNickle, 2023; Sica et al., 2024; Zhong et al., 2024). However, it remains to be seen whether these fluctuations can be linked to the benefits and costs of each infection state under specific ecological conditions.

The prevalence of Totivirus-infected yeast isolates is a greater challenge to explain. Yeast with totivirus infections, but lacking the satellite, are expected to incur the costs of maintaining the totivirus without gaining the toxin-encoding benefits provided by the satellite (Boynton, 2019; Chau et al., 2023a). Whilst intracellular selection should favour mutant totiviruses with higher virulence, selection should act against yeast with virulent totiviruses because of their reduced fitness compared to those infected by benign totiviruses. In low population density environments, the loss of the satellite may be inconsequential. However, at high population densities, when toxins are most effective as interference competition strategies (Greig & Travisano, 2008), yeast with totivirus infections should be selected against. Thus, the persistence of this infection-based polymorphism is likely to depend on density dependent processes, in which their persistence relies on low densities and minimal competition. The near-complete replacement of yeast with totivirus infections by those without infections in 2021 may be explained by the costs of totiviruses to their yeast hosts – a burden uninfected yeast do not experience.

We hypothesised that infection states would diverge alongside their host genotypes, given that *S. cerevisiae* primarily reproduces asexually, thereby linking infection state directly to host lineages. For this pattern not to emerge, frequent sexual reproduction would need to occur between genotypes with differing infection states. Such events would mix evolutionarily isolated lineages, homogenize infections and/or decouple infection states from host lineages, ultimately disrupting the expected pattern of genotype clustering by infection state that arises from clonal reproduction. The DAPC results demonstrate that, with high mean successful assignment (MSA), we can cluster host genotypes by their infection states. This supports our hypothesis that totiviruses and the satellites primarily experience vertical transmission from parent to progeny, because genotypes that are most closely related to each other host the same infection states. This was most evident when genotype frequencies were included (Figure 3a) and genotypes with infection state transitions are excluded (Figure 3c).

Under a stepwise infection transition scenario, the totivirus-infected state acts as an intermediate transition state between being coinfected with both the totivirus and its satellite, and being infection-free. Our DAPC models support this scenario (Figure 3), given that the totivirus-infection genotype cluster tended to fall within an intermediate genotype space between the coinfected and infection-free clusters. A stepwise infection state transition scenario is also supported by genotypes that were sampled across years (Figure 2). While resampled genotypes across years most frequently maintained their infection state, when they did experience infection state transitions, it was almost exclusively in a stepwise manner (Figure 2). Though rare, these transitions demonstrate that yeast are losing their totiviruses, and that satellites are also being lost from those where a coinfection was present, within natural populations across generations. Removal of the satellite nucleic acid under laboratory conditions is considerably more straightforward than that of the totivirus, which we find moderate support for in this study (Figure 2; Gao et al., 2019; Wickner, 1974). The mechanisms by which *S. cerevisiae* eliminates the totivirus are largely unknown, though various mechanisms have been identified for controlling their copy number proliferation (Gao et al., 2019; Rowley et al., 2016).

Our transition matrix indicates that, statistically, genotypes in the natural population can acquire totiviruses and their associated satellites (i.e., infection gains; Figure 2). The only known mechanism by which these elements can be acquired is through cytoplasmic mixing during sexual reproduction. If sexual reproduction were responsible for infection gains, we would expect recombination to disrupt the combinations of microsatellite alleles that define genotypes. As a result, the same genotypes would be unlikely to reappear across years, since recombination rarely produces identical configurations across multiple diploid loci. Consequently, genotypes with higher infection states would not be expected to belong to the same genetic lineages as those with matching microsatellite profiles from previous years. Furthermore, by emphasizing inter-genotypic relationships (Figure 3c)—through clone correction and the exclusion of genotypes with multiple infection states—a distinct clustering by infection state becomes apparent. Such a pattern would be unlikely if frequent sexual reproduction between genotypes with different infection states were occurring. While the infection-state-driven structuring suggests that, if sexual events do occur, they are largely confined to genotypes sharing the same infection status, overall, sex is likely to be infrequent and not responsible for the statistical acquisition of infections over time.

Intragenotypic gains of totiviruses and/or satellites across years may indicate that individuals with higher infection states were already present in prior years but went undetected, indicating an issue of the sampling effort. Our results are, however, also consistent with—and suggestive of—the possibility of extracellular acquisition of totiviruses and/or their satellites, given that genotypes can statistically acquire mycoviruses over time despite the lack of evidence for sexual reproduction. While this phenomenon has not been previously detected and is considered unlikely—because mycoviruses do not cause cell lysis (Ghabrial et al., 2015; Ghabrial & Suzuki, 2009) – the idea has gained support in recent years, and several mechanisms have been proposed (Buivydaite et al., 2024). This is not to imply that our results provide evidence for the extracellular acquisition of mycoviruses; rather, they are more consistent with extracellular acquisition being the mechanism for mycovirus gains, rather than sexual transmission. Given the growing interest in the possibility of extracellular acquisition of mycoviruses (Buivydaite et al., 2024), experimental approaches should be employed to test the feasibility of this phenomenon.

Our transition probability matrix demonstrates that genotypes experience transitions in infection state between sampling years. This is particularly perplexing, given our DAPC clustering results, because over multiple years this process should disrupt the clustering of host genotypes by infection status. One explanation for this could be that host genotypes have optimal infection states, whereby offspring with transitioned infection states are less fit than the original infection states over longer periods of time. There is extensive literature on host genes affecting the maintenance of dsRNAs in *S. cerevisiae* (Magliani et al., 1997; Wickner, 1996a, 1996b). Recessive mutations in a number of *mak* genes have been shown to result in the loss of the satellite during haploid stages (Magliani et al., 1997). In many cases, the totivirus remains unaffected by these recessive mutants whilst the satellite is eliminated (*MAK4, MAK5, MAK6, MAK7, MAK14* and *MAK15*) (Magliani et al., 1997; Wickner & Leibowitz, 1976b, 1979b). Three genes (*MAK3, MAK10* and *PET18)* are known to be necessary for the maintenance of totiviruses as well as their satellites (Fujimura et al., 1986; Fujimura & Wickner, 1987; Schmitt & Tipper, 1992; Tercero et al., 1993; Tercero & Wickner, 1992). Tracking variants in genes involved in infection maintenance could provide insight into the causes of transitions, as well as the factors influencing their persistence or suboptimality.

We report the persistence of infection-based competitive polymorphism in a natural population of *S. cerevisiae.* We demonstrate that infection state divergence occurs with host genotypes, and that transitions in infection type are occurring by result of intracellular conflict, and perhaps also by sexual reproduction. However, transitions to suboptimal infection types are unlikely to persist over longer periods as they would disrupt the clustering of host genotypes by infection types. The results of our study underscore the need for more resolute longitudinal studies, both intra-seasonal and inter-annual, to investigate the dynamics of infection-based competitive polymorphism in the killer yeast system. Such studies should integrate host genomics, infection dynamics, and both competition and fitness assays.

## Acknowledgements

This work was supported by ETH Research Grant ETH-23 20-1.

## Supplementary Materials

**Table S1.**
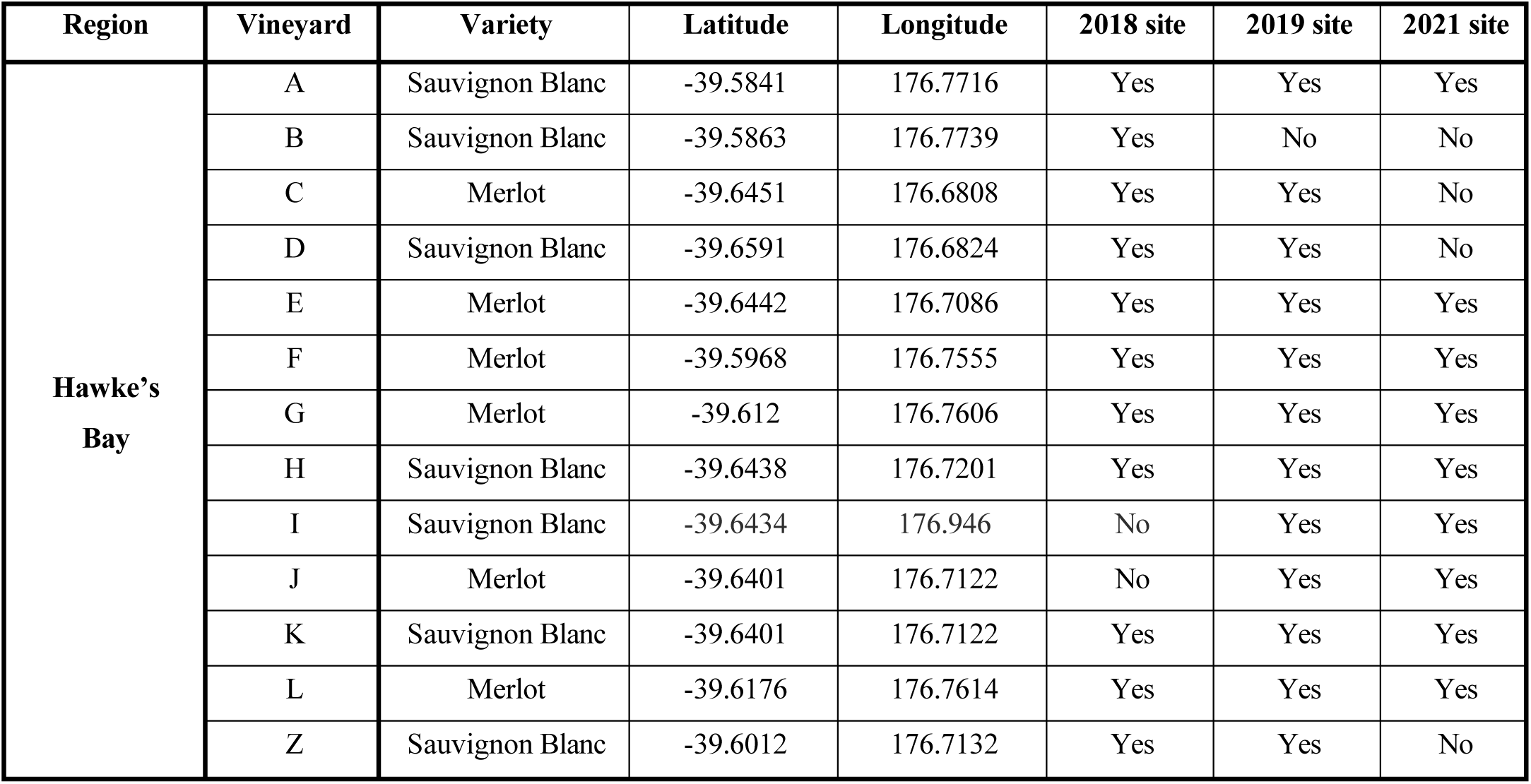
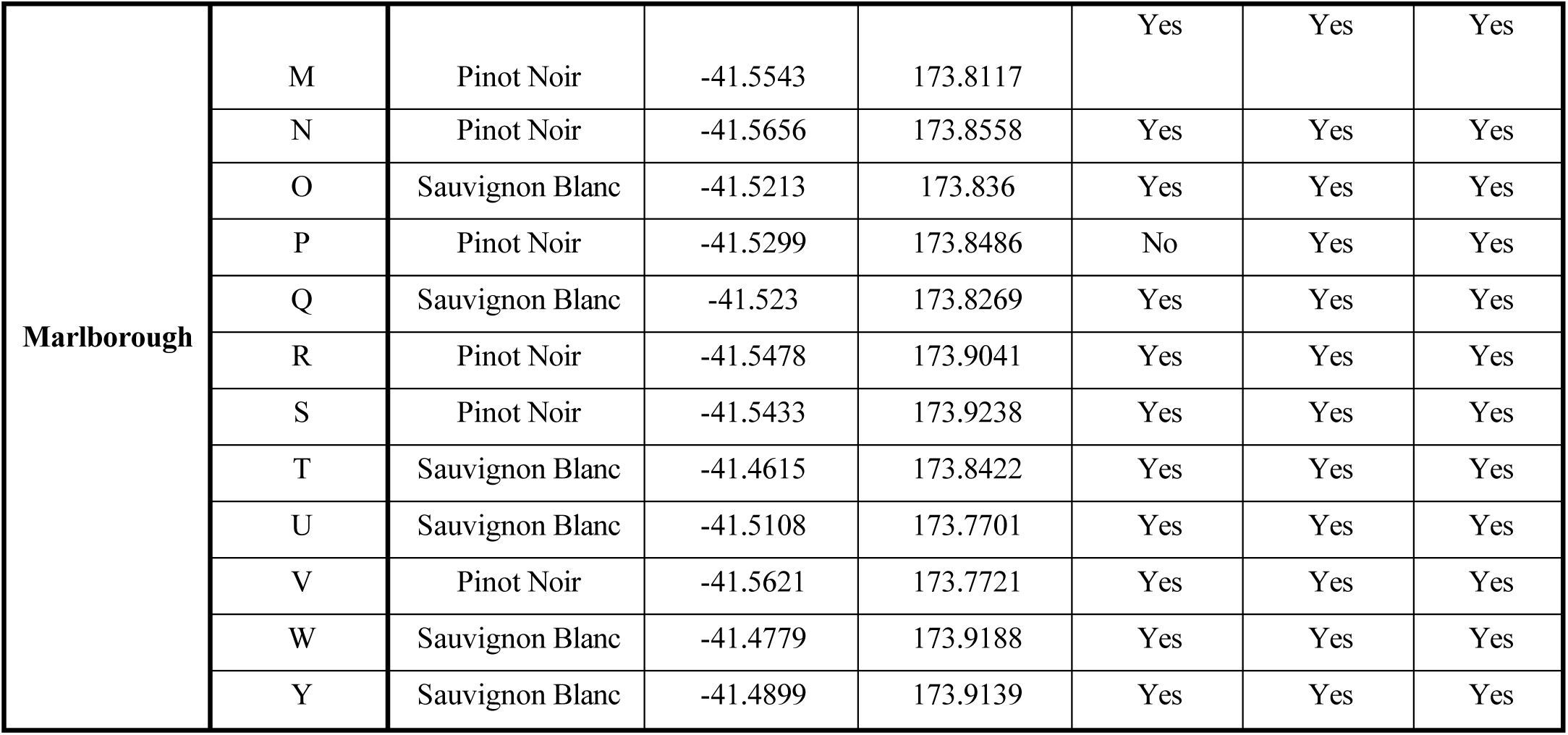
Information on the vineyards from which *Saccharomyces cerevisiae* was sampled.

| Region | Vineyard | Variety | Latitude | Longitude | 2018 site | 2019 site | 2021 site |
| --- | --- | --- | --- | --- | --- | --- | --- |
| Hawke's Bay | A | Sauvignon Blanc | -39.5841 | 176.7716 | Yes | Yes | Yes |
|  | B | Sauvignon Blanc | -39.5863 | 176.7739 | Yes | No | No |
|  | C | Merlot | -39.6451 | 176.6808 | Yes | Yes | No |
|  | D | Sauvignon Blanc | -39.6591 | 176.6824 | Yes | Yes | No |
|  | E | Merlot | -39.6442 | 176.7086 | Yes | Yes | Yes |
|  | F | Merlot | -39.5968 | 176.7555 | Yes | Yes | Yes |
|  | G | Merlot | -39.612 | 176.7606 | Yes | Yes | Yes |
|  | H | Sauvignon Blanc | -39.6438 | 176.7201 | Yes | Yes | Yes |
|  | I | Sauvignon Blanc | -39.6434 | 176.946 | No | Yes | Yes |
|  | J | Merlot | -39.6401 | 176.7122 | No | Yes | Yes |
|  | K | Sauvignon Blanc | -39.6401 | 176.7122 | Yes | Yes | Yes |
|  | L | Merlot | -39.6176 | 176.7614 | Yes | Yes | Yes |
|  | Z | Sauvignon Blanc | -39.6012 | 176.7132 | Yes | Yes | No |
| **Marlborough** | M | Pinot Noir | -41.5543 | 173.8117 | Yes | Yes | Yes |
|  | N | Pinot Noir | -41.5656 | 173.8558 | Yes | Yes | Yes |
|  | O | Sauvignon Blanc | -41.5213 | 173.836 | Yes | Yes | Yes |
|  | P | Pinot Noir | -41.5299 | 173.8486 | No | Yes | Yes |
|  | Q | Sauvignon Blanc | -41.523 | 173.8269 | Yes | Yes | Yes |
|  | R | Pinot Noir | -41.5478 | 173.9041 | Yes | Yes | Yes |
|  | S | Pinot Noir | -41.5433 | 173.9238 | Yes | Yes | Yes |
|  | T | Sauvignon Blanc | -41.4615 | 173.8422 | Yes | Yes | Yes |
|  | U | Sauvignon Blanc | -41.5108 | 173.7701 | Yes | Yes | Yes |
|  | V | Pinot Noir | -41.5621 | 173.7721 | Yes | Yes | Yes |
|  | W | Sauvignon Blanc | -41.4779 | 173.9188 | Yes | Yes | Yes |
|  | Y | Sauvignon Blanc | -41.4899 | 173.9139 | Yes | Yes | Yes |

**Table S2.**
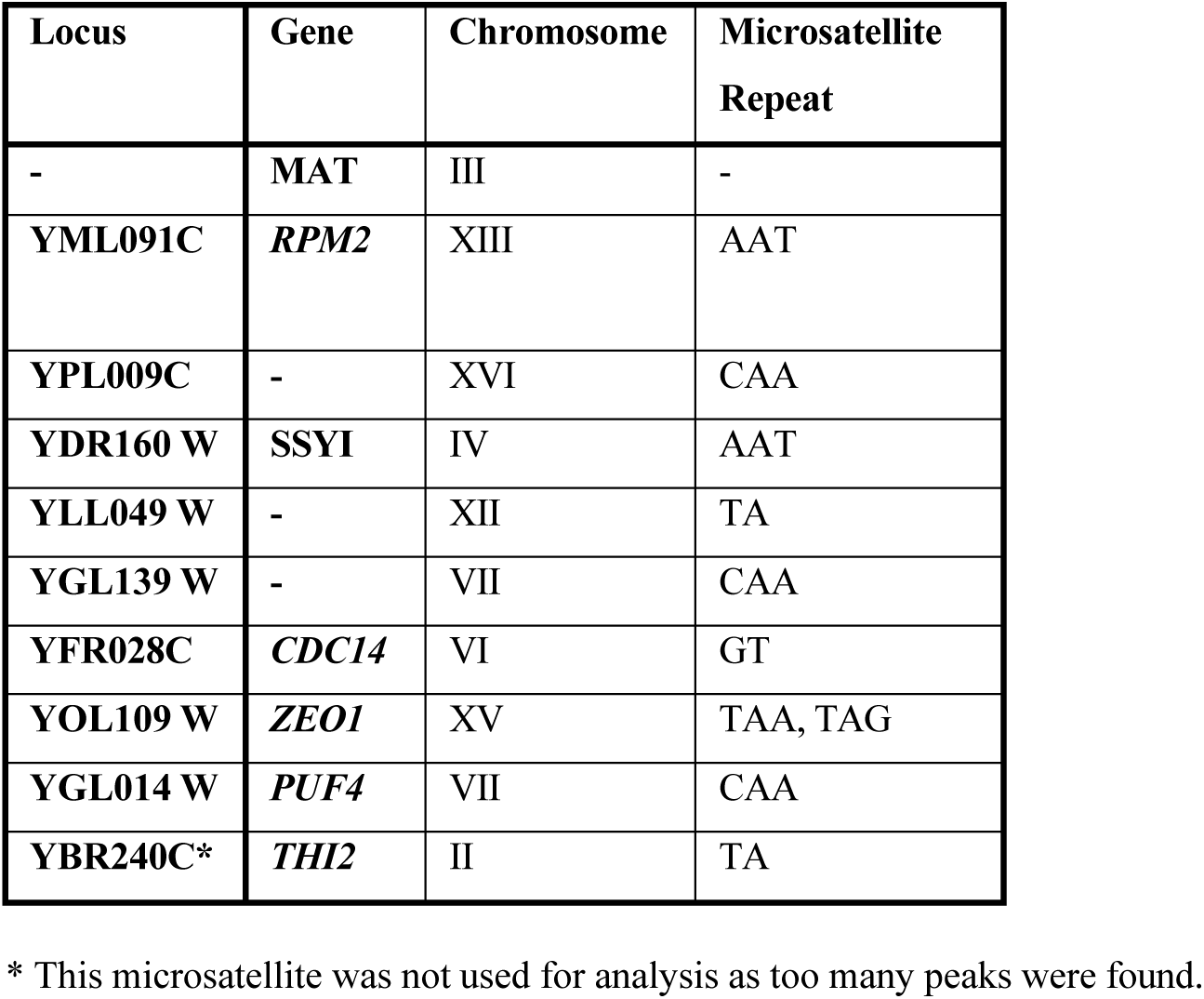
Microsatellite marker information.

**Image S1.**
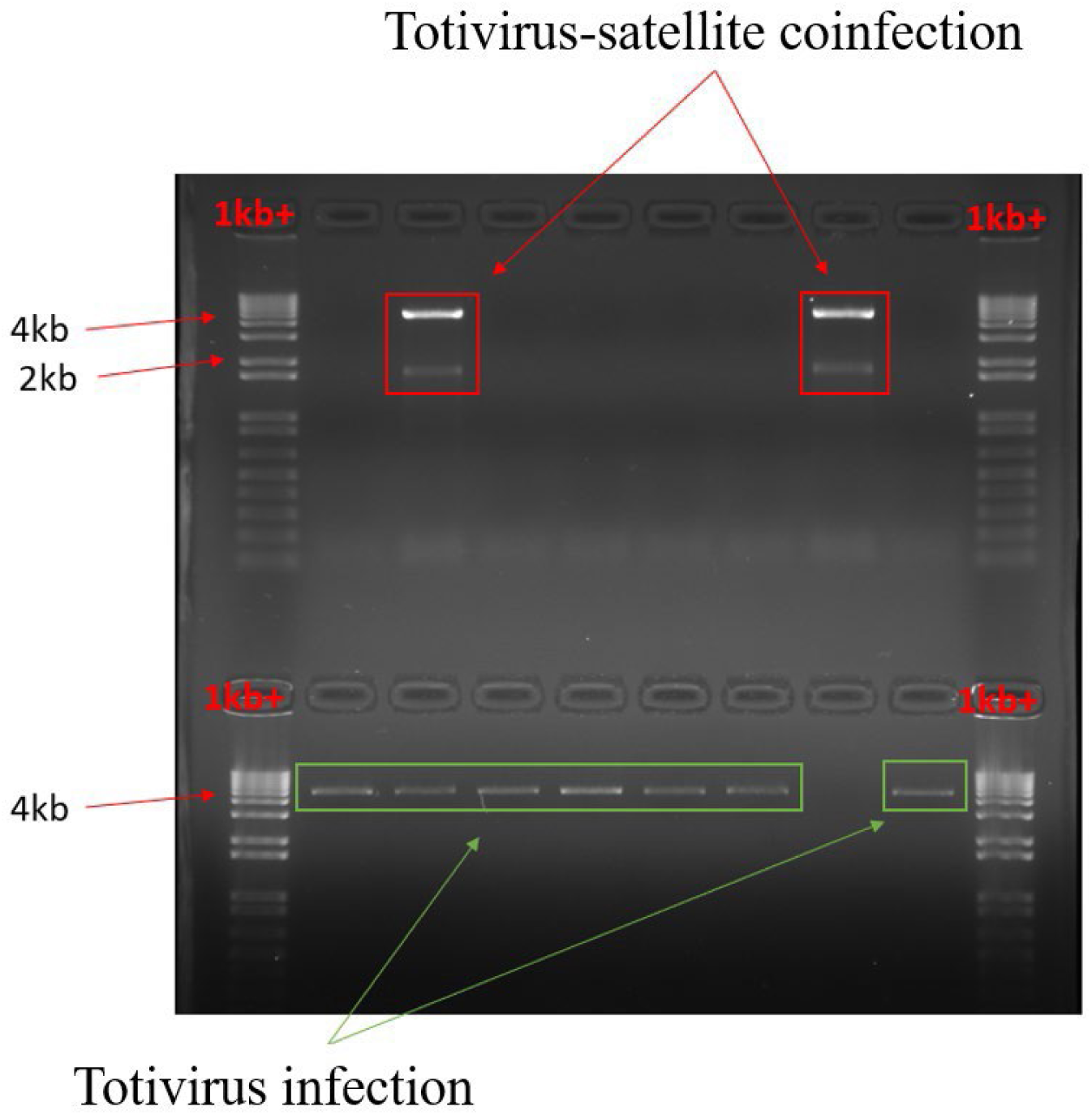
Photographic example of an agarose gel with all the three virus profiles represented.

**Figure S1:**
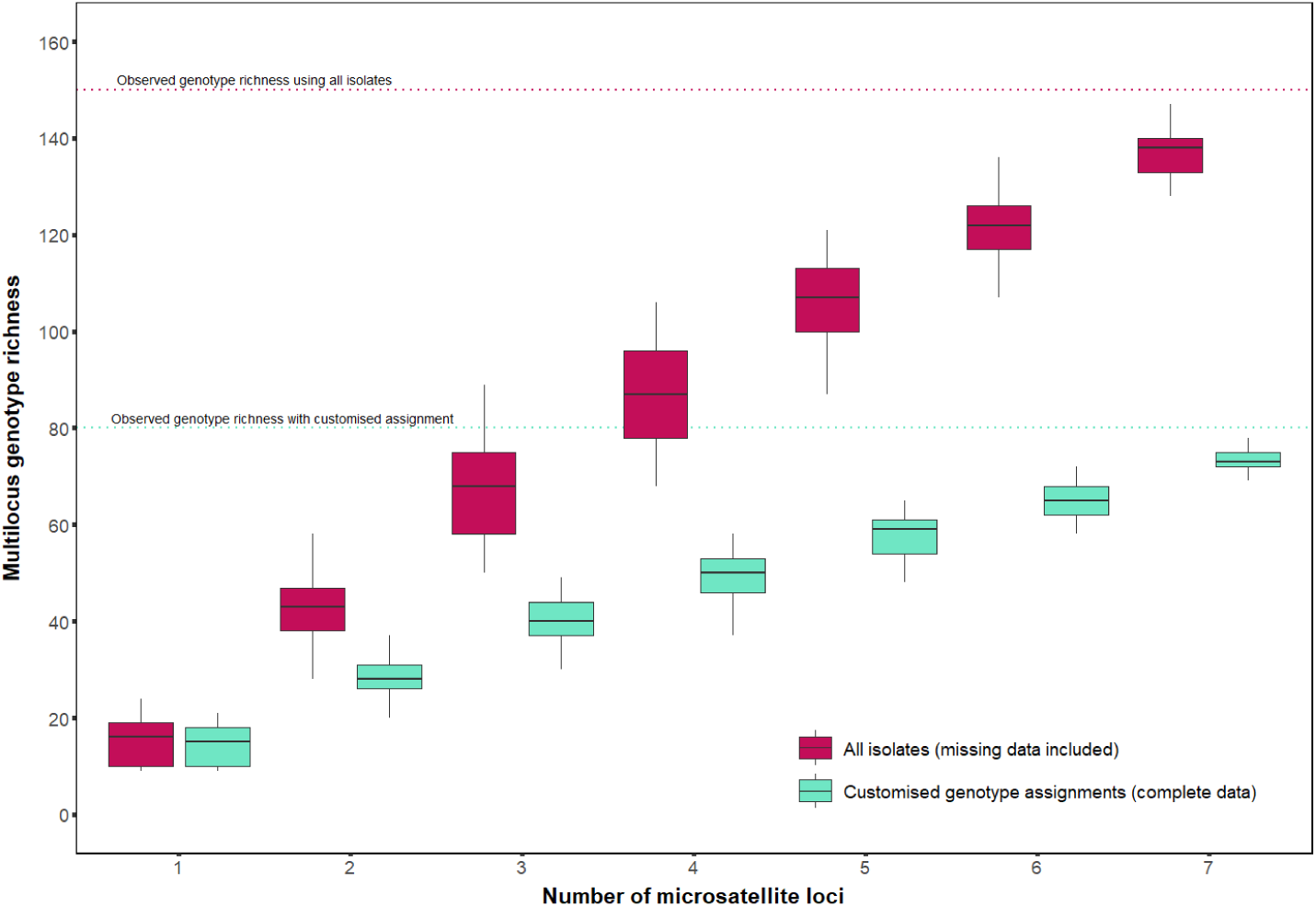
Multilocus genotype accumulation for all isolates including those with missing data (red) and customised genotypes whereby genotypes with missing data were nested into other genotypes with more complete data or excluded if they could be nested into multiple other genotypes (pale blue). Horizontal lines indicate the observed genotype richness for each genotype accumulation curve.

**Figure S2:**
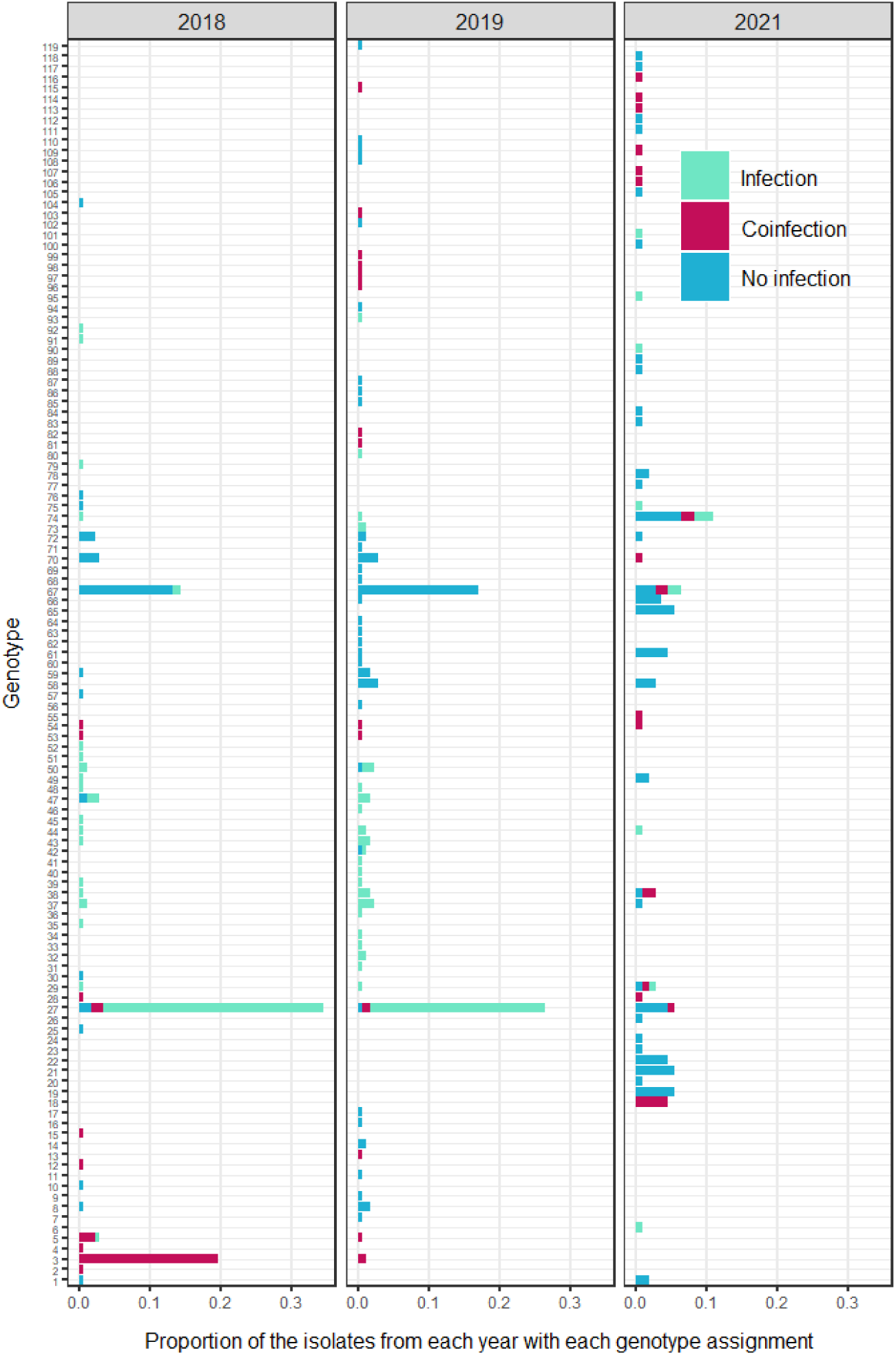
The proportions of each unique genotype per year across the three years.

